# Growth-Aware Optogenetic Control for Cross-Scale Disturbance Rejection in Bacterial Batch Cultures

**DOI:** 10.64898/2026.04.04.716495

**Authors:** Hari R. Namboothiri, Chelsea Y. Hu

## Abstract

Population-level disturbances can disrupt molecular gene expression through growth-dependent host–circuit coupling, creating a cross-scale challenge for synthetic biology. Here, we use a multiscale model to develop growth-aware optogenetic control strategies that maintain a target molecular-scale gene expression signal following population-level dilution in bacterial batch cultures. Simulations predict distinct disturbance-response regimes within a model that couples population growth to intracellular dilution and expression capacity. At intermediate disturbance magnitudes, growth-dependent feedback gains reduce signal overshoot relative to the fixed-gain controller tested. At large disturbances, additional population-based feedforward compensation accelerates signal recovery. Mapping performance across disturbance magnitudes and times identifies when each architecture is beneficial, with the predicted rankings retained under tested parameter perturbations. These results establish model-guided design rules for cross-scale disturbance rejection, showing how population-level information can inform feedback and feedforward regulation of molecular gene expression.

---

Precise regulation of gene expression is a key objective in synthetic biology, with applications ranging from living therapeutics to biomanufacturing^1,2^. However, disturbances and control objectives can arise at different biological scales. In bacterial batch cultures, population-level perturbations such as culture dilution can alter molecular gene-expression dynamics through growth-dependent changes in intracellular dilution and expression capacity. Maintaining a molecular expression target therefore requires rejecting disturbances that propagate across scales. This coupling creates a distinct control challenge: a population perturbation can change both the expression output and the dynamics governing its recovery.

Optogenetic cell-silicon systems provide a means to address this challenge by combining light-actuated gene expression with real-time measurements and computer-based control^3-5^. Previous studies have demonstrated setpoint tracking and disturbance rejection using proportional–integral (PI), proportional–integral–derivative (PID), and model predictive controllers^6-10^. Much of this work uses continuous-culture or microfluidic platforms that maintain approximately steady growth conditions. In batch culture, growth transitions continually reshape gene-expression dynamics, complicating controller design and the interpretation of disturbance responses^11,12^. The resulting question is how population-state information should inform molecular-output regulation when disturbances enter through the growth dynamics.

Our previous work established the Gene Expression Across Growth Stages (GEAGS) framework to connect population growth with molecular circuit dynamics^11^,^13^. We subsequently combined growth-aware modeling with the LED Embedded Microplate for Optogenetic Studies (LEMOS) platform to demonstrate molecular-scale setpoint tracking in batch cultures^4^. Here, we extend this foundation to cross-scale disturbance rejection: maintaining a gene-expression target following a population-level perturbation. The coupled model provides a basis for tracing how the disturbance affects molecular expression and identifying appropriate corrective action.

Using simulated culture dilutions, we identify how disturbance magnitude and growth state change molecular recovery trajectories and the relative performance of different control architectures. We develop growth-aware feedback and population-based feedforward strategies and compare their performance across disturbance magnitudes and times. The simulations show that growth-dependent feedback gains suppress overshoot at intermediate disturbances, whereas additional feedforward compensation accelerates recovery after large disturbances. These findings provide model-guided design rules for using information at the population scale to regulate gene expression at the molecular scale.

We studied cross-scale disturbance rejection in an optogenetic system whose molecular expression dynamics are coupled to population growth. The CcaSR two-component system regulates sfGFP expression in response to green and red light (Fig. 1A)^14^. We used the published GEAGS framework and its growth-dependent rate-modifier functions to couple population growth to molecular expression dynamics (Fig. 1B)^4^. The model formulation and biological basis of these functions are described in the original studies; the parameters and equations used here are provided in the SI section 1. Simulations reproduce the measurement and actuation sequence of LEMOS, with fluorescence per cell (*P*_*m*_) as the controlled output and light duty cycle as the manipulated input (Fig. 1C)^4^. The output *P*_*m*_, provides a population-averaged readout of molecular expression.

**Figure 1.**
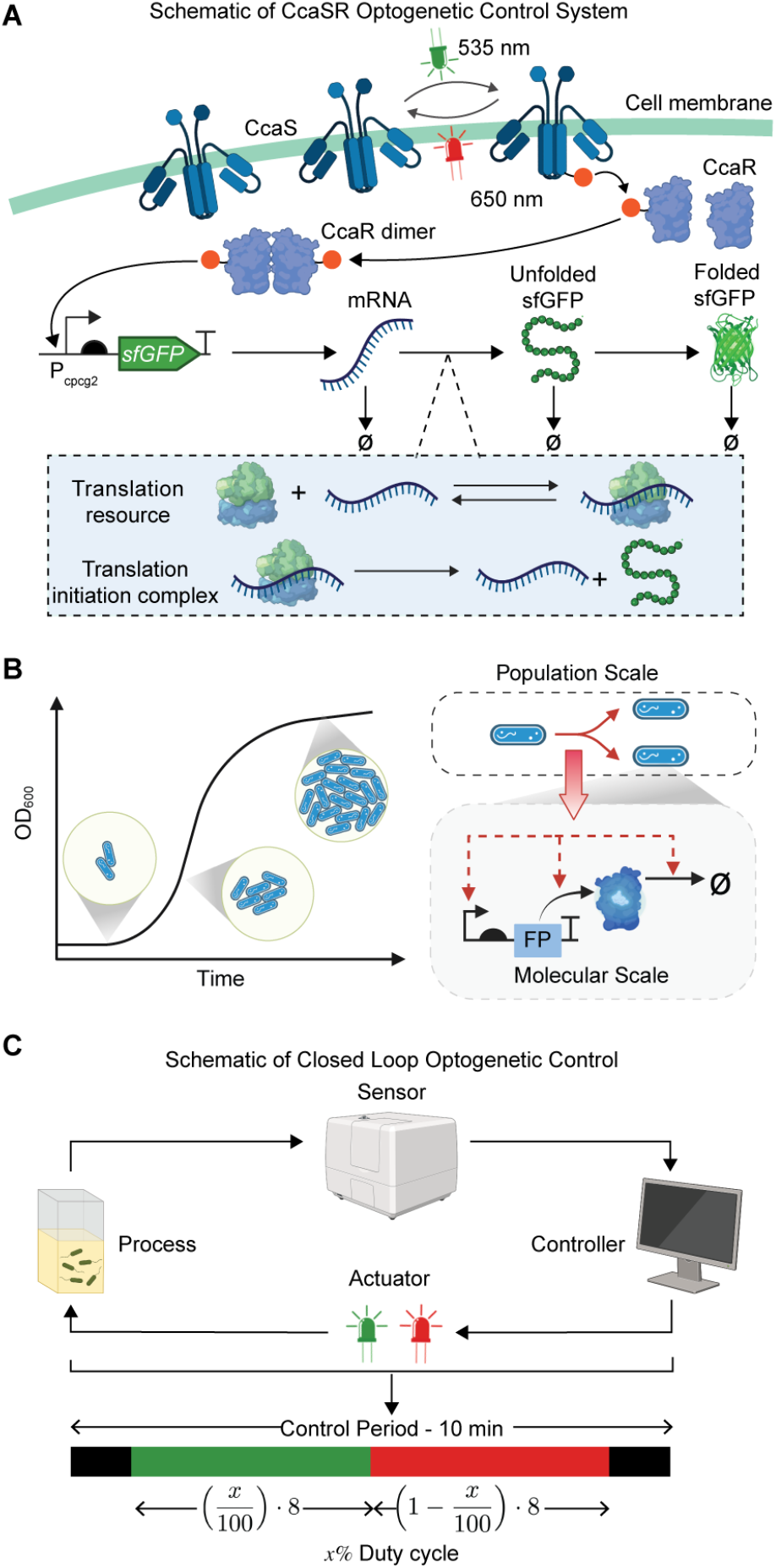
Multiscale model formulation and system description. (**A**) Schematic of the CcaSR two-component optogenetic system. Green light (535 nm) activates CcaS/CcaR signaling and induces PcpcG2-sfGFP expression, whereas red light (650 nm) represses expression. The molecular species and interactions shown correspond to the chemical reactions represented in the dynamic model. (**B**) Schematic illustrating how population-scale dynamics influence the molecular scale gene expression dynamics. (**C**) Schematic of the closed loop optogenetic control system. The system operates on a 10-minute control period, with the LEDs OFF during the first and last minute. The duty cycle sets the fraction of the remaining 8-minute actuation window with green light ON, with red light ON for the rest.

Within this coupled system, we asked whether molecular expression could be maintained at a target level following a population-level disturbance. We introduced simulated culture dilutions by reducing cell density at a specified time while leaving the expression setpoint unchanged (Fig. 2A). In the GEAGS model, normalized cell density determines the magnitude of the growth-dependent rate modifiers; a density reduction therefore changes the molecular reaction rates governing the subsequent expression trajectory. This formulation captures a cross-scale disturbance pathway: the perturbation enters at the population level, while its downstream effects must be rejected at the molecular output (Fig. 2A). We used this framework to examine how disturbance magnitude and growth state shape the expression response and the corrective action required.

**Figure 2.**
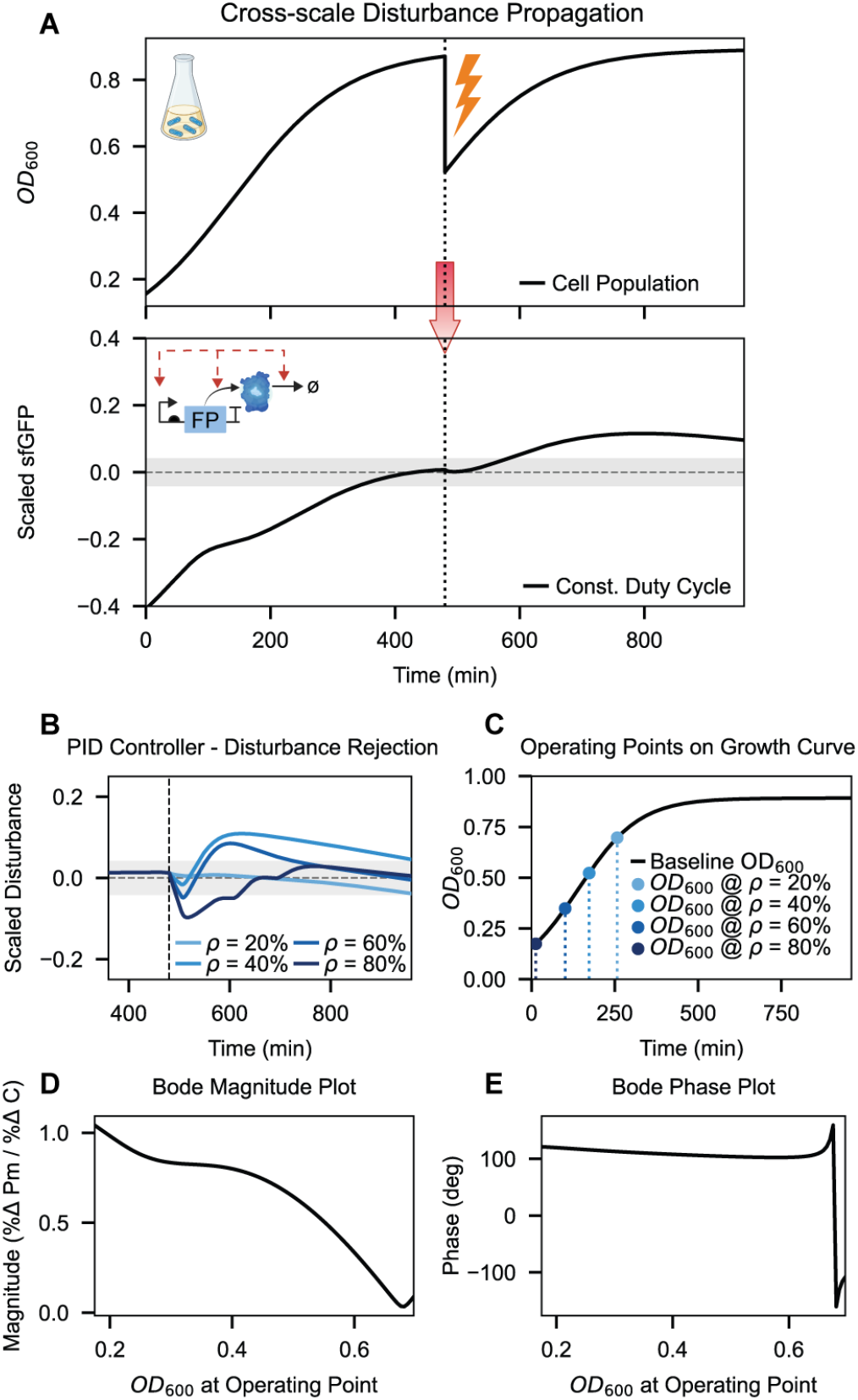
Closed-loop disturbance rejection and open-loop frequency response analysis. (**A**) Schematic illustrating the propagation of population-scale disturbance into molecular-scale gene expression dynamics. Upper panel demonstrates the disturbance in bacterial population caused by dilution at a fixed time point and the bottom panel illustrates the resulting open-loop dynamics of sfGFP expression under constant duty cycle. (**B**) Fixed-gain PID controller response under varying disturbance percentages. Grey shaded region indicates a 5% tolerance band around the setpoint. (**C**) Operating points along the growth curve. (**D**) Bode magnitude plot illustrating the decrease in system gain as OD increases.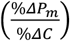 represents the amplitude ratio of the percentage deviation of fluorescence per cell (*P*_*m*_) to a unit-amplitude sinusoidal perturbation in population state (*C*). (**E**) Bode phase plot demonstrating a pronounced increase in temporal lag at higher OD_600_. The phase represents the temporal lag between the oscillations in *C* and in the *P*_*m*_ response.

Population-level dilution altered the molecular expression trajectory even when illumination remained constant (Fig. 2A), demonstrating disturbance propagation through the growth– expression coupling represented in GEAGS. Because the model links population state to both intracellular dilution and expression capacity, a density reduction changes multiple processes governing fluorescence per cell. The resulting response therefore depends on the balance of these processes. We examined how this coupled response affects regulation by the fixed-gain PID controller from our previous setpoint-tracking study.

Population-level dilution altered the molecular expression trajectory even when illumination remained constant (Fig. 2A), demonstrating disturbance propagation through the growth– expression coupling represented in GEAGS. Because the model links population state to both intracellular dilution and expression capacity, a density reduction changes multiple processes governing fluorescence per cell, including transcription, translation, mRNA degradation and protein degradation rates^11,15-18^. The resulting response therefore depends on the balance of these processes. We examined how this coupled response affects regulation by the fixed-gain PID controller from our previous setpoint-tracking study.

To determine how these effects influence closed-loop regulation, we applied the fixed-gain PID controller from our previous setpoint-tracking study^4^ and introduced density reductions of *ρ* = 20– 80% at 480 min (Fig. 2B). A 20% reduction produced little deviation in *P*_*m*_ because the culture remained close to stationary phase. Intermediate reductions of 40% and 60% shifted the population into a range of higher expression capacity and were followed by transient overshoot. The overshoot was greater after the 40% reduction than after the 60% reduction, demonstrating that a larger population disturbance did not necessarily produce a larger transient increase in expression. By contrast, an 80% reduction produced a more pronounced initial decrease in *P*_*m*_, consistent with stronger growth-associated dilution, followed by recovery under feedback control.

To understand why the fixed-gain PID controller produced different responses across disturbance conditions, we next examined how population state shapes the underlying gene-expression dynamics. Using the GEAGS model, we evaluated representative growth states corresponding to the population densities reached after different dilution conditions (Fig. 2C) and quantified how small, periodic changes in population density affect fluorescence per cell across different timescales (see Methods). These responses were visualized using Bode plots, which describe both the strength of the molecular response and the delay between a population perturbation and the resulting change in fluorescence. The analysis showed that cultures at lower OD_600_ were more sensitive to population perturbations, whereas the response weakened as cultures approached stationary phase. Phase changed little across most of the OD_600_ range, then shifted sharply as the response magnitude approached zero. (Fig. 2D–E). Together with the disturbance trajectories in Fig. 2B, these results indicate that the molecular scale consequences of population scale perturbations, and the recovery action required, vary with growth state.

These responses distinguish two control demands within the same coupled system: suppressing the expression overshoot associated with intermediate dilutions and accelerating recovery from the initial expression loss associated with large dilutions. The fixed-gain PID controller exhibited different recovery trajectories across these conditions, motivating controller designs that use population-state information to adjust corrective action. We therefore examined whether growth-dependent feedback gains could reduce overshoot and whether additional population-based feedforward compensation could improve recovery after large disturbances.

To address these different recovery demands, we introduced a growth-aware adaptive PID controller with online gain scaling based on normalized population density 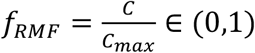,. Here, adaptation refers to updating controller gains at each control interval according to a predefined rule based on the current simulated population density. We prescribed linear relationships between normalized population density and both the proportional gain and derivative time constant, reducing feedback aggressiveness at lower densities and increasing it as the population recovers. The controller takes the form

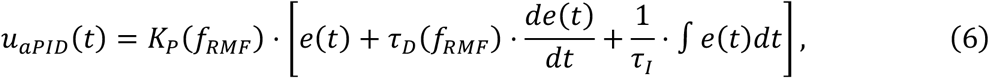

where *u*_*aPID*_ (*t*) is the feedback controller output, *K*_*P*_(*f*_*RMF*_)=*K*_*P*_· *f*_*RMF*_ +*K*_*P*,0_,τ_*D*_(*f*_*RMF*_)= τ_*D*_· *f*_*RMF*_+ τ_*D*,0_, and τ_*I*_ are the effective proportional gain, derivative time constant, and integral time constant respectively, and *e*(*t*) = setpoint™*P*_*m*_ is the tracking error. The basal terms *K*_*P*,0_ and τ_*D*,0_ maintain feedback action at low population density. Following dilution, the lower value of *f*_*RMF*_ reduces feedback aggressiveness; as the population recovers, the gains increase. We evaluated whether this prescribed growth-dependent adjustment could suppress the overshoot observed with fixed-gain control.

Growth-aware feedback reduced overshoot following intermediate dilutions (Fig. 3B-C). The improvement was particularly pronounced after a 60% density reduction, where the transient increase in expression observed with the fixed-gain controller was largely eliminated. However, the same gain reduction slowed signal recovery after an 80% dilution, when the dominant initial response was a loss of fluorescence per cell (Fig. 3D). At 20% dilution, the controllers produced similar trajectories (Fig. 3A). These simulations reveal a tradeoff: reducing feedback aggressiveness limits overshoot under intermediate disturbances but weakens the corrective action needed after large disturbances. Growth-dependent feedback therefore addresses one consequence of cross-scale disturbance propagation while motivating an additional mechanism for accelerating recovery from substantial expression loss.

**Figure 3.**
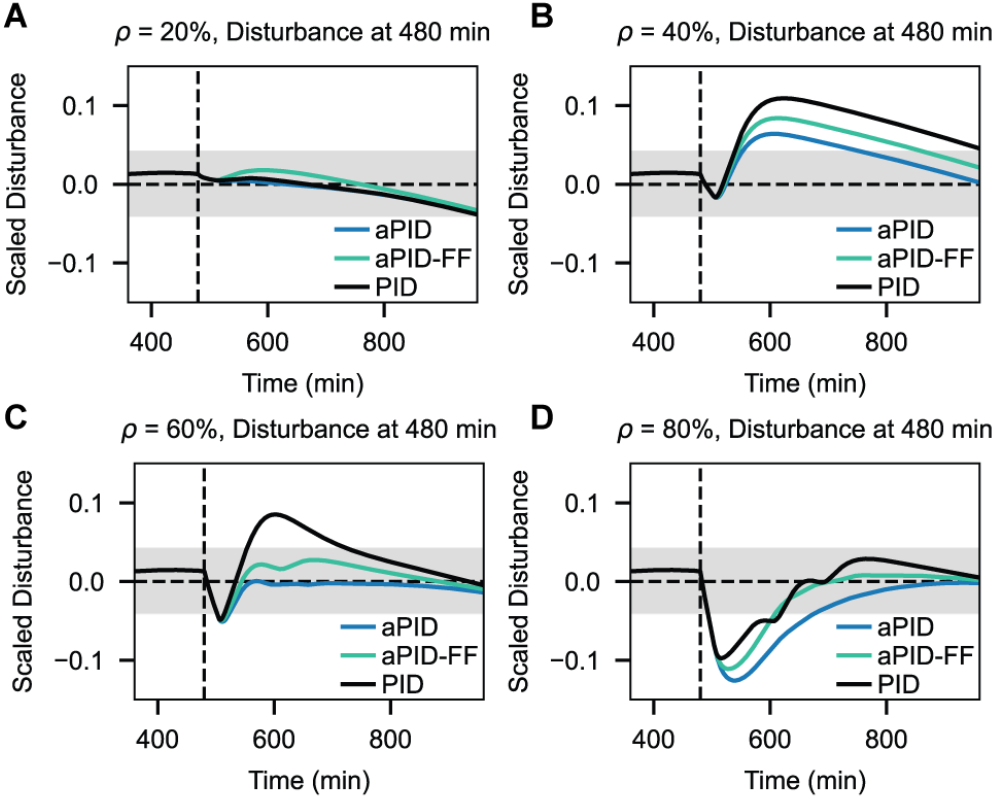
Comparison of the adaptive controllers with fixed-gain PID controller under varying disturbance percentages. The grey shaded region represents the 5% tolerance band.

To accelerate recovery after large dilutions, we supplemented growth-aware feedback with a feedforward term based on the population disturbance. Whereas feedback control responds to deviations in fluorescence per cell, population-based feedforward uses the upstream density change to initiate corrective action before its full molecular consequence develops. In the simulations, the feedforward term uses the current population density. An experimental implementation would obtain the corresponding population-state information from OD_600_ measurement. Because population growth evolves independently of the light input in this model, population density can serve as an upstream signal to anticipate and compensate for disturbances in molecular gene expression. We defined the feedforward input as

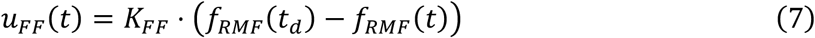

where *f*_RMF_ (*t*_d_) is the normalized population density immediately before dilution and *K*_*FF*_ is the feedforward gain selected via manual tuning using GEAGS simulations. The feedforward term is inactive before the disturbance. After dilution, its contribution scales with the density change relative to the pre-disturbance state. The combined control input is

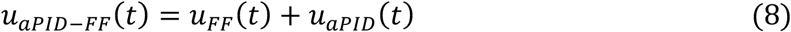

This architecture uses population information in two complementary ways: to adjust feedback gains as growth conditions change and to provide a direct corrective input following a population disturbance.

After an 80% dilution, feedforward compensation accelerated recovery relative to growth-aware feedback alone, while retaining lower overshoot than the fixed-gain controller (Fig. 3D). Its benefit was less pronounced at intermediate dilutions, where the combined controller produced slightly greater overshoot than growth-aware feedback alone (Fig. 3B-C). This behavior is consistent with a feedforward contribution that persists while the population remains below its pre-disturbance density, even after molecular expression has recovered to the setpoint. At 20% dilution, adding feedforward produced little change in the response (Fig. 3A). These results show that population-based feedforward addresses the recovery limitation of growth-aware feedback after large disturbances but can introduce excess corrective action when overshoot suppression is the primary control demand.

To identify when each control architecture is beneficial, we compared their performance across a (7X7) grid of dilution magnitudes, ranging from 20% to 80%, and disturbance times, ranging from 360 to 540 min. Disturbance-rejection performance was quantified using the integral of time-weighted absolute error (ITAE):

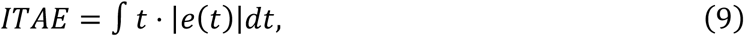

which penalizes persistent deviations from the molecular expression setpoint. We plotted its inverse,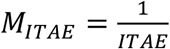, so that higher values indicate better performance (Fig. 4A–D). This analysis extended the representative trajectories into a map of controller performance across population perturbations applied at different stages of growth.

**Figure 4.**
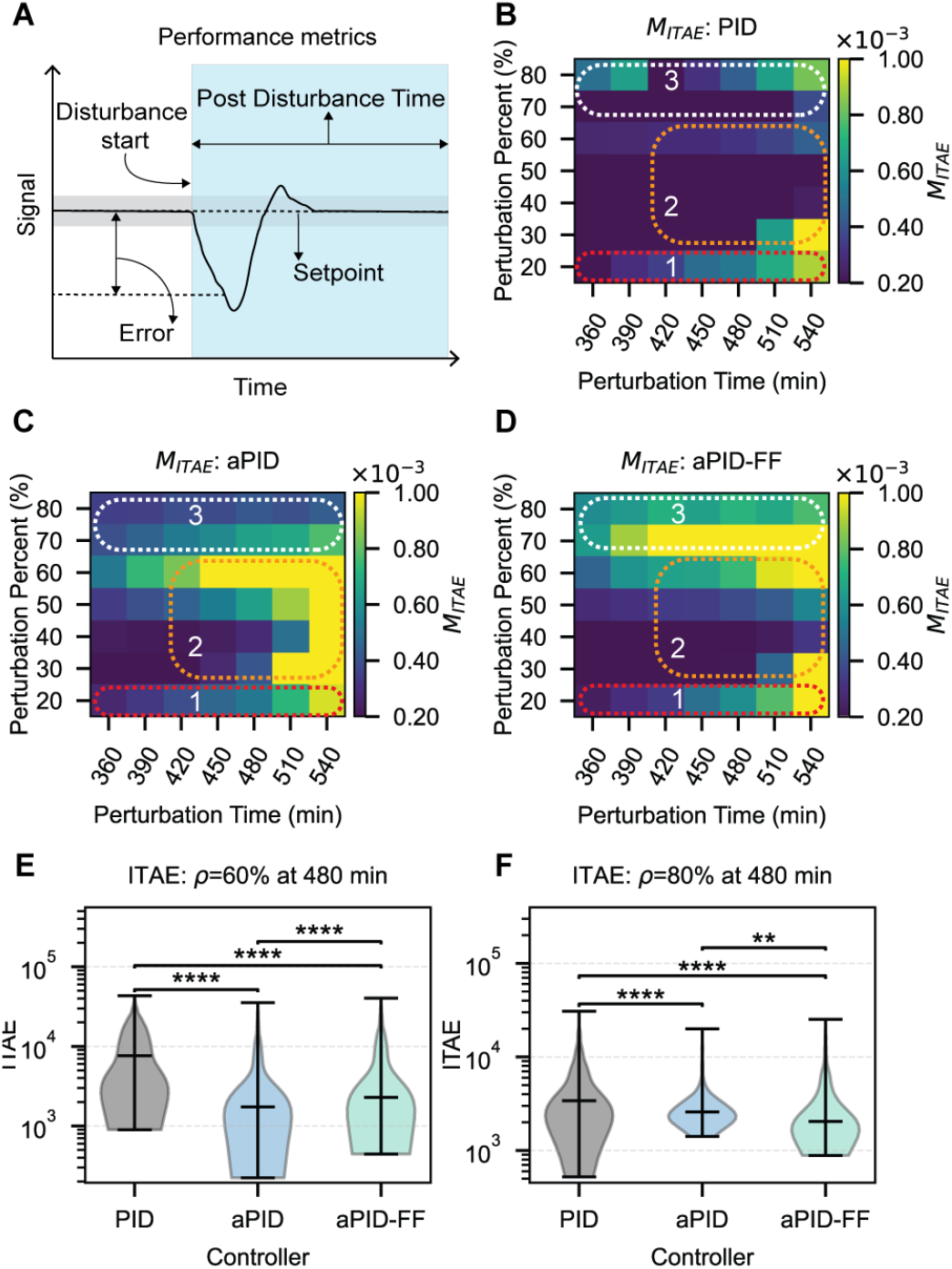
Comparison of the performance and robustness of the three controllers across disturbance conditions. (**A**) Schematic of the performance metrics used to calculate the combined metric *M*. (**B-D**) Heatmaps showing the variation in *M*_*ITAE*_ across perturbation times and perturbation percents for each of the three controllers. The dashed ellipses indicate the different operating regimes outlined in the text. (**E-F**) Violin plots showing distribution of ITAE across the Monte Carlo trajectories. The upper and lower bars represent the maximum and minimum value of ITAE, and the center bar represents the median. The asterisks denote statistical significance between the two groups, determined by paired Student’s t-tests.

The simulations identified three operating regimes. At small dilutions, the controllers performed similarly, indicating limited benefit from growth-dependent gains or additional feedforward compensation. At intermediate dilutions applied in the late growth stages, growth-aware feedback generally performed best, consistent with the importance of suppressing the expression overshoot associated with increased gene expression capacity. At large dilutions, the combined feedback–feedforward controller achieved the best performance across the disturbance times tested. In this regime, the population-based corrective input accelerated recovery from the initial expression loss, while growth-aware feedback limited subsequent overshoot. Thus, the preferred architecture depended on the molecular recovery demand created by the population disturbance.

We next examined whether the controller rankings persisted under model-parameter uncertainties. For representative intermediate and large dilutions, we evaluated 1,000 Monte Carlo trajectories with parameters varied by ±15% from their nominal values after disturbance onset (see Methods). At a 60% dilution applied at 480 min, growth-aware feedback retained the lowest median ITAE; at an 80% dilution applied at the same time, the combined feedback–feedforward controller retained the lowest median ITAE (Fig. 4E–F). These results support the predicted rankings under the tested parameter variations.

Together, the comparisons provide model-guided controller-selection rules: growth-dependent feedback is favored when overshoot suppression dominates, whereas additional population-based feedforward is favored when rapid recovery from expression loss is required. These rules describe the conditions evaluated here. Their practical use depends on how population perturbations alter growth and expression dynamics in the culture being controlled.

This study extends growth-aware modeling and molecular-scale setpoint tracking to cross-scale disturbance rejection. Population-state information serves both to adjust feedback gains and to drive feedforward compensation for disturbances whose consequences emerge in molecular gene expression. The simulations identify distinct benefits of these strategies: growth-dependent feedback suppresses overshoot after intermediate dilutions, whereas population-based feedforward accelerates recovery after large dilutions. These findings provide model-based design rules that connect controller selection to the competing effects of growth-dependent dilution and expression capacity.

These design rules are supported by parameter-perturbation analyses within the growth– molecular coupling represented in GEAGS, where simulated dilution immediately changes growth-dependent reaction rates through normalized population density. Their applicability to systems with physiological memory or delayed adjustment to fresh medium remains to be tested. Experimental evaluation under different dilution conditions would clarify how these dynamics shape controller performance and when population-state information offers the greatest benefit.

## Methods

### Frequency Response Analysis

Open-loop frequency-response analysis was performed by linearizing the GEAGS model at each operating point shown in Fig. 2C, yielding a local linear time-invariant (LTI) state-space representation. Sinusoidal perturbations in population state were defined as the input, and the corresponding percentage deviation in fluorescence per cell was defined as the output. Bode plots were then used to characterize the magnitude and phase of this input–output response across perturbation frequencies and growth states. At each operating point, the Bode magnitude and phase were computed using the *scipy*.*signal*.*bode* function. To relate the frequency-response analysis to the timescale of the simulated dilution response, we focused on the low-frequency regime and 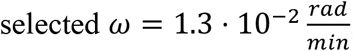, as a representative frequency, corresponding to a period of 480 min. This timescale captures the evolution of the system following dilution over the remaining simulation window. The state-space formulation and detailed derivation of the frequency-response analysis are provided in SI Section 2.

### Parameter Uncertainty and Robustness Analysis

Controller robustness was evaluated using 1,000 Monte Carlo simulations for each representative disturbance condition. All kinetic parameters in the GEAGS model were varied independently and simultaneously within ±15% of their nominal values using uniform sampling, while logistic-growth and controller parameters were held constant. Each trajectory used one sampled parameter set, applied from the disturbance onset onward so that all simulations shared the same pre-disturbance initial conditions and controller history (Fig. S1). Controller performance was quantified using the ITAE, calculated over the post-dilution period by numerically integrating the time-weighted absolute setpoint tracking error using the trapezoidal rule.

## Acknowledgment

The authors would like to thank Ayush Pandey (UC Merced) for their insightful discussions. This research is supported by funding provided to C.Y.H. by Texas A&M Engineering Experiment Station (TEES).

## Code Availability

All the codes needed to evaluate and reproduce the results in the paper are available at the GitHub repository: https://github.com/synbiosystems/Growth-Aware-Disturbance-Rejection-Controllers-LEMOS.git

