## Supplementary Information for "Growth-Aware Optogenetic Control for Cross-Scale Disturbance Rejection in Bacterial Batch Cultures"

### 1 GEAGS Model Formulation

#### *1.1 Model Description and Equations*

The signal sensing and transduction dynamics of the CcaSR system is represented by the chemical reaction network (CRN)[1]:

$$CcaS \underset{k_{rvrs}}{\overset{k_{fwd}}{\rightleftarrows}} CcaS_P \tag{S1}$$

$$CcaS_P + CcaR \underset{k_{R_u}}{\overset{k_{R_b}}{\rightleftarrows}} CcaS + CcaR_P \tag{S2}$$

$$CcaR_p + CcaR_p \underset{k_{Rp_u}}{\overset{k_{Rp_b}}{\rightleftarrows}} Ac \tag{S3}$$

Under green light, $CcaS$ autophosphorylates at the full rate $k_{fwd} = k_{green} + k_{fb}$; in its absence, only the basal rate applies ($k_{fwd} = k_{fb}$). Under red light, $CcaS_p$ dephosphorylates at the full rate $k_{rvrs} = k_{red} + k_{rb}$; in its absence only the basal rate ($k_{rvrs} = k_{rb}$) applies. $CcaR$ undergoes reversible phosphorylation with forward and reverse rate constants $k_{R_b}$ and $k_{R_u}$ respectively. Similarly, $CcaR_p$ reversibly forms the dimer $Ac$, with forward and reverse rate constants $k_{Rp_b}$ and $k_{Rp_u}$ respectively. The dynamics of the signaling pathway is modeled as:

$$\begin{aligned}\frac{dCcaS}{dt} &= -\left(k_{green} + k_{fb}\right) \cdot CcaS + \left(k_{red} + k_{rb}\right) \cdot CcaS_p \\ &\quad + k_{R_b} \cdot CcaS_p \cdot CcaR - k_{R_u} \cdot CcaS \cdot CcaR_p\end{aligned} \tag{S4}$$

$$\begin{aligned}\frac{dCcaS_p}{dt} &= \left(k_{green} + k_{fb}\right) \cdot CcaS - \left(k_{red} + k_{rb}\right) \cdot CcaS_p \\ &\quad - k_{R_b} \cdot CcaS_p \cdot CcaR + k_{R_u} \cdot CcaS \cdot CcaR_p\end{aligned} \tag{S5}$$

$$\frac{dCcaR}{dt} = -k_{R_b} \cdot CcaS_p \cdot CcaR + k_{R_u} \cdot CcaS \cdot CcaR_p \tag{S6}$$

$$\begin{aligned}\frac{dCcaR_p}{dt} &= k_{R_b} \cdot CcaS_p \cdot CcaR - k_{R_u} \cdot CcaS \cdot CcaR_p \\ &\quad - k_{R\square\square_b} \cdot CcaR_p^2 + k_{Rp_u} \cdot Ac\end{aligned} \tag{S7}$$

$$\frac{dAc}{dt} = k_{Rp_b} \cdot CcaR_p^2 - k_{Rp_u} \cdot Ac \tag{S8}$$

In closed batch cultures, growth rate varies continuously, so conventional single-scale models with constant-growth assumptions cannot capture the shifting physiological state of cells. The GEAGS framework resolves this limitation by coupling intracellular gene expression to population-level logistic growth [2]:

$$\frac{dC}{dt} = C \cdot k_{gr} \cdot \left(1 - \frac{C}{C_{\max}}\right), \tag{S9}$$

where $C$ is the cell population, $k_{gr}$ is the logistic growth rate, and $C_{\max}$ is the carrying capacity, defined as the maximum population size bounded by the nutrient and space constraints. To link molecular-scale dynamics with population-scale growth, GEAGS uses custom rate modifier functions (RMFs), which define empirical relationships between growth-dependent reactions and bacterial growth phases. Each RMF is a function of the variable $f_{RMF} = \frac{C}{C_{max}} \in (0,1)$, which indicates the proximity of the population to the carrying capacity:

$$\alpha = 1 - f_{RMF} \tag{S10}$$

$$\delta = \frac{f_{RMF}^n}{1 + f_{RMF}^n} \tag{S11}$$

$$\gamma = [f_{RMF} \cdot (1 - f_{RMF})]^m \tag{S12}$$

The RMF $\alpha$ (Eq. S10), quantifies how the logistic growth rate changes in batch culture as the population approaches carrying capacity. In the model, $\alpha$ modulates the dilution rates of molecular species and reduction in growth dependent mRNA degradation. The RMF $\delta$ (Eq. S11), represents the effect of growth-phase transitions on key intracellular processes, increasing sharply as cells enter stationary phase. In the model, $\delta$ captures the upregulation of sfGFP degradation during this transition. The RMF $\gamma$ (Eq. S12), models the growth phase-dependent molecular process efficiency, peaking in mid-log phase when growth is fastest. In the model, $\gamma$ modulates the transcription rate, sfGFP maturation rate and the coarse-grained translation resource ($R$) availability.

Although gene translation is inherently a higher-order process, it is often simplified as a first-order reaction dependent solely on mRNA concentration, assuming abundant translational resources. Here, we explicitly model the growth-dependent availability of translational resources to account for resource competition. This refinement is important because mRNA levels differ substantially under green and red light, which likely alters the availability of translational resources and leads to different translation rates in the two conditions[3]. The multiscale gene expression model is described as:

$$\frac{dM}{dt} = \beta'_m \cdot \left(\frac{Ac}{Kc + Ac} + l_0\right) - (d'_m + d'_{dil}) \cdot M - k_{tli_b} \cdot R_{free} \cdot M + k_{tli_u} \cdot C_{tic} + k_{tl} \cdot C_{tic} \tag{S13}$$

$$\frac{dC_{tic}}{dt} = k_{tli_b} \cdot R_{free} \cdot M - k_{tli_u} \cdot C_{tic} - k_{tl} \cdot C_{tic} \tag{S14}$$

$$\frac{dP}{dt} = k_{tl} \cdot C_{tic} - \left(d'_p + d'_{dil}\right) \cdot P - k'_{fold} \cdot P \tag{S15}$$

$$\frac{dP_m}{dt} = k'_{fold} \cdot P - \left(d'_p + d'_{dil}\right) \cdot P_m \tag{S16}$$

$$\frac{dC}{dt} = k_{gr} \cdot C \cdot \left(1 - \frac{C}{C_{max}}\right) \tag{S17}$$

Where, the modified growth-dependent rate equations are:

$$\beta'_m = \beta_m \cdot \gamma \tag{S18}$$
$$d'_m = d_m \cdot \alpha \tag{S19}$$
$$d'_p = d_p \cdot \delta \tag{S20}$$
$$d'_{dil} = d_{dil} \cdot \alpha \tag{S21}$$
$$k'_{fold} = k_{fold} \cdot (\gamma + b_{fold}) \tag{S22}$$

Here $M, C_{TIC}, P,$ and $P_m$ denote mRNA, translation initiation complex, unfolded sfGFP and matured sfGFP respectively. $\beta_m$ is the transcription rate; $K_c$ is the promoter activation coefficient; $l_0$ is the leak coefficient; $k_{tl}$ is the translation rate; $d_m$ and $d_p$ are the mRNA and protein degradation rates respectively; $k_{fold}$ and $b_{fold}$ are the maturation, and basal maturation rates of sfGFP; and $d_{dil}$ is the dilution rate due to cell growth. In addition, $k_{tli_b}$ is the binding rate of freely available translation resource ($R_{free}$) and mRNA, and $k_{tli_u}$ is the unbinding rate of the coarse-grained translation initiation complex $C_{tic}$. The dynamics of $R$ and $C_{tic}$ are described as:

$$R_{total} = R_{max} \cdot \gamma \tag{S23}$$
$$R_{free} = R_{total} - C_{TIC} \tag{S24}$$

#### *1.2 Model Species and Parameters*

Model species

**Table S1** GEAGS model species

| Species | Description |
|---|---|
| $M$ | mRNA coding for sfGFP |
| $P$ | Unfolded sfGFP |
| $P_m$ | Folded sfGFP |
| $CcaS$ | CcaS (membrane protein) |
| $CcaS_P$ | Phosphorylated CcaS |
| $CcaR$ | CcaR (response regulator protein) |
| $CcaR_P$ | Phosphorylated CcaR |
| $Ac$ | Transcription activation complex |
| $C_{tic}$ | Translation initiation complex |
| $R$ | Coarse-grained translation resource |
| $C$ | Cell population |

Model parameters

**Table S2** GEAGS model parameters

| Parameter | Description | Unit | Estimate |
|---|---|---|---|
| $\beta_m$ | Transcription rate per plasmid | $nM \cdot min^{-1}$ | 2.8e1 |
| $l_0$ | Leak coefficient of promoter | $N/A$ | 1e-5 |

| | | | |
|---|---|---|---|
| $K_c$ | Dissociation constant of Ac binding to promoter | $nM$ | 4.5e1 |
| $d_m$ | mRNA degradation rate constant | $min^{-1}$ | 2.7e-1 |
| $k_{tli_b}$ | $C_{tic}$ formation rate | $nM^{-1} \cdot min^{-1}$ | 4e1 |
| $k_{tli_u}$ | $C_{tic}$ dissociation rate | $min^{-1}$ | 1e1 |
| $k_{tl}$ | Translation elongation rate | $min^{-1}$ | 2 |
| $d_p$ | Protein degradation rate | $min^{-1}$ | 8e-4 |
| $k_{fold}$ | sfGFP maturation rate | $min^{-1}$ | 3e-1 |
| $b_{fold}$ | Basal coefficient for $k_{fold}$ | $N/A$ | 1 |
| $k_{green}$ | Phosphorylation rate of CcaS under green light | $min^{-1}$ | 8e-1 |
| $k_{fb}$ | Basal phosphorylation rate of CcaS | $min^{-1}$ | 3e-2 |
| $k_{red}$ | Dephosphorylation rate of CcaS under red light | $min^{-1}$ | 1.3 |
| $k_{rb}$ | Basal dephosphorylation rate of CcaS under red light exposure | $min^{-1}$ | 8e-1 |
| $k_{R_b}$ | Phosphorylation rate of CcaR by $CcaS_p$ | $nM \cdot min^{-1}$ | 5e-2 |
| $k_{R_u}$ | Dephosphorylation rate of $CcaR_p$ | $nM \cdot min^{-1}$ | 2.5e1 |
| $k_{R_{pb}}$ | Forward dimerization rate of $CcaR_p$ | $nM \cdot min^{-1}$ | 5.5e1 |
| $k_{R_{pu}}$ | Reverse dimerization rate of $CcaR_p$ | $min^{-1}$ | 8e-2 |
| $R_{max}$ | Max. total R availability | $nM$ | 4 |
| $n$ | Exponent of $\gamma$ | $N/A$ | 8.9e-1 |
| $n_{delta}$ | Hill coefficient of $\delta$ | $N/A$ | 5.5 |
| $C_0$ | Initial condition for cell population | $counts$ | 4.69e7 |
| $C_{max}$ | Max. cell population (holding capacity) | $counts$ | 7.14e8 |
| $k_{gr}$ | Logistic growth rate | $min^{-1}$ | 1.1e-2 |

### 2 Frequency Response Analysis

#### 2.1 Nonlinear Model Representation:

The GEAGS model is described by $n$ coupled ODEs encoding the mass-balance kinetics of all the molecular species as well as the population species:

$$\dot{x} = f(x) \tag{S25}$$

where $x = [x_1, \dots, x_n] \in \mathbb{R}^n$ is the state vector of species concentrations (including $C$ and $P_m$), and $f: \mathbb{R}^n \to \mathbb{R}^n$ maps the nonlinear dynamics. The baseline trajectory $x^*(t)$ is obtained by numerical integration of Eqs. (S1 – S24) from the initial condition $x_0$.

#### 2.2 Operating Point Selection:

The frequency response is evaluated at an array of operating points mapped by the disturbance magnitude $\rho \in [20\%, 80\%]$. Denoting the cell population at the disturbance time as $C_{ref} = C^*(t_d)$, the post-disturbance population is

$$C_\rho = C_{ref} \cdot \left(1 - \frac{\rho}{100}\right). \tag{S26}$$

The operating-point time is the time at which the baseline trajectory satisfies the equation $C^*(t_\rho) = C_\rho$ (Fig. 2C, main text), and the corresponding operating point is $x_{op}(\rho) = x^*(t_\rho)$.

#### 2.3 Local Linearization:

To linearize the nonlinear model at each operating point $x_{op}(\rho)$, we estimate the Jacobian of $f$ numerically using fourth-order central-difference. For each state index $j$, an adaptive perturbation step

$$h_j = \max\left(10^{-8}, 10^{-2} \cdot \left|x_{op,j}\right|\right) \tag{S27}$$

is applied, and the $(i, j)$-th entry of the Jacobian matrix $A$ is

$$A = \left.\frac{\partial f_i}{\partial x_j}\right|_{t_\rho} = \frac{-f_i(x_i + 2h_j) + 8f_i(x_i + h_j)}{12h_j} - \frac{8f_i(x_i - h_j) - f_i(x_i - 2h_j)}{12h_j}. \tag{S28}$$

Because $C$ is treated as an external input, it is removed from the state vector, and we define a reduced state vector $\tilde{x} \in \mathbb{R}^{\tilde{n}}$ ($\tilde{n} = n - 1$), which contains all the remaining species[4]. Partitioning $A$ yields the reduced system matrix $A_{red} \in \mathbb{R}^{\tilde{n} x \tilde{n}}$ representing the linearized dynamics of $\tilde{x}$, and the input vector $B = \frac{\partial \tilde{f}}{\partial C} \in \mathbb{R}^{\tilde{n}}$, quantifying the influence of $C$ on those dynamics.

*2.4 State Space Representation:*

We define a scalar input $u$ to represent a sinusoidal percentage perturbation in $C$ relative to the local operating point population $C_\rho = C^*(t_\rho)$. A unit-amplitude input corresponds to a physical perturbation in $C$ as

$$\partial C = -\left(\frac{C_\rho}{100}\right) \cdot u, \tag{S29}$$

where the negative sign reflects the convention that $u > 0$ denotes a reduction in $C$. Substituting the expression of $\partial C$ into the linearized dynamics gives the modified input matrix

$$B' = -\left(\frac{C_\rho}{100}\right) \cdot \frac{\partial \tilde{f}}{\partial C}\bigg|_{x_{op}} \in \mathbb{R}^{\tilde{n}} \tag{S30}$$

The output, $P_m$, is expressed as a percentage deviation from its operating-point value $P_{m,op}$, yielding the output matrix

$$C_{mat} = \left(\frac{100}{P_{m,op}}\right) \cdot e_y \in \mathbb{R}^{\tilde{n}} \tag{S31}$$

where $e_y$ selects the $P_m$ entry from $\tilde{x}$. The feedthrough matrix is set to be $D = \mathbf{0}$. Using the matrices defined in Eqs. (S28 – S31), we can define the continuous linear time invariant (LTI) state-space model in terms of deviation variables as

$$\partial \dot{\tilde{x}} = A_{red} \partial \tilde{x} + B' u, \tag{S32}$$

$$\partial y = C_{mat} \partial \tilde{x} \tag{S33}$$

and the corresponding scalar transfer function as

$$G(s) = C_{mat}(sI - A_{red})^{-1} B'. \tag{S34}$$

### 3 Robustness Analysis

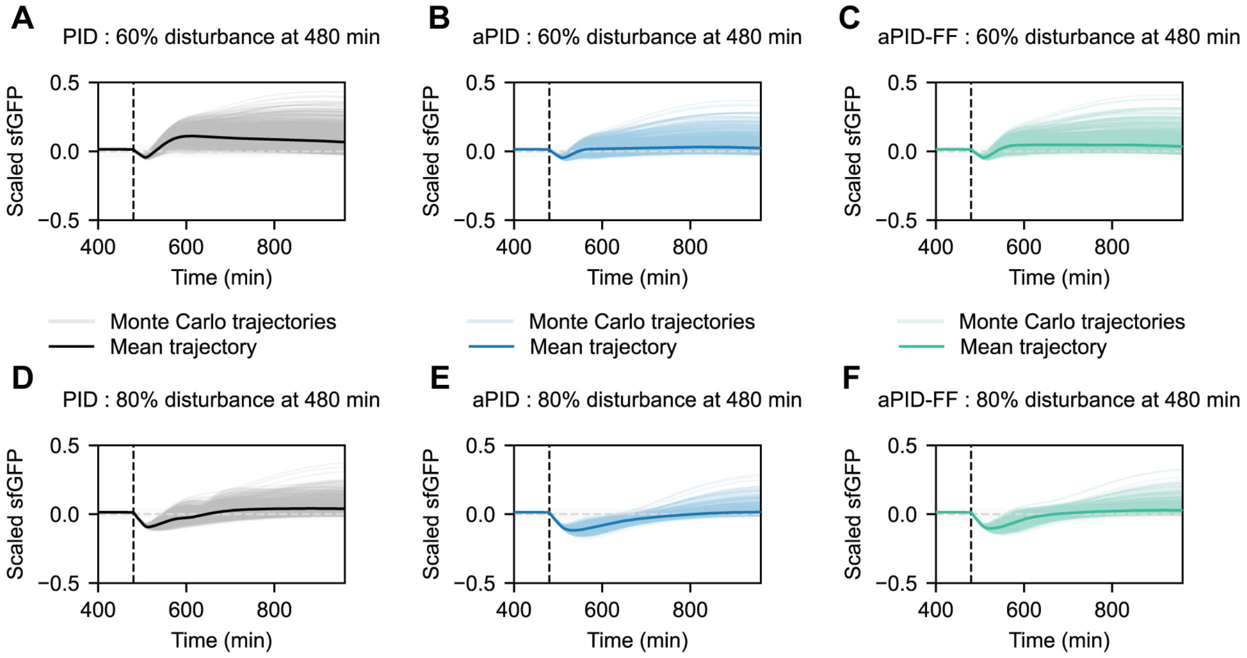


**Figure S1 Robustness analysis of disturbance rejection controllers.** Random trajectories (n=1000) were generated by uniformly sampling parameters about ±15% of their nominal values.

4 Åström, K. J. & Murray, R. *Feedback systems: an introduction for scientists and engineers*. (Princeton university press, 2021).
